# Targeted editing of pericentromeric satellite DNA alters sensitivity to meiotic drive

**DOI:** 10.64898/2026.01.18.700156

**Authors:** D.G. Eickbush, J. Rahmat, M. Lindsay, J. Bowers, N.J. Fuda, A.M. Larracuente

## Abstract

Eukaryotic genomes are abundant in satellite DNA (satDNA): large blocks of tandemly-repeated sequences that accumulate in heterochromatic genome regions. SatDNAs are dynamic in their genomic location and abundance across species. Some satDNAs overlap essential genome regions such as centromeres and telomeres, but even pericentromeric satDNA can have effects on phenotypes, raising questions about their functional significance. However, it remains unclear whether these effects depend on satDNA sequence, copy number, higher-order structural organization, or genomic context. The highly repetitive nature of satDNA arrays has long hindered detailed genomic and genetic analyses. Recent advances in long-read sequencing now facilitate both the detailed characterization of satDNA structure and the development of more targeted approaches to genetic analysis. Here we present a sequential CRISPR/Cas9-based strategy to make mutations in satDNA arrays and demonstrate its utility using an autosomal pericentromeric satDNA in *Drosophila melanogaster* called *Responder (Rsp). Rsp* is the target of a sperm-killing male meiotic driver, *Segregation Distorter* (*SD*), where sensitivity to sperm killing positively correlates with *Rsp* copy number. Using our CRISPR/Cas9 approach, we generated an allelic series of *Rsp* deletion and expansion variants in two genetic backgrounds and examined their effects on spermatogenesis. Our approach produced precise satDNA variants efficiently, with minimal detectable off-target effects. The resulting mutations affect sensitivity to *SD* that scale with *Rsp* copy number. This work establishes a new framework for experimentally dissecting satDNA function and provides insights into the evolutionary and functional roles of satDNA in genome organization.

## Introduction

Satellite DNAs (satDNA) are highly repetitive non-coding genetic elements that accumulate in eukaryotic genomes, especially around centromeres, telomeres, and on sex chromosomes. SatDNAs consist of arrays of tandem repeats and can occupy hundreds of kilobases or megabases of sequence, typically packaged as heterochromatin. The misregulation of satellites is associated with age-related phenotypes and diseases like cancer^1-4^, and fertility defects (reviewed in ^5^), although the cause of these phenotypes is poorly understood. Some satDNAs can play roles in chromosome organization and segregation, although it is unclear which properties of satellites are important for these roles (e.g. structural or sequence^6,7^). SatDNAs help coordinate essential nuclear organization through satellite-binding proteins ^8^ and accumulate in genome regions that have essential roles like centromeres. Genome regions rich in satDNA are generally prone to instability at the sequence, chromatin, and transcription levels. SatDNAs are among the most rapidly evolving genome regions across species. These dynamics can have consequences for genome evolution over long evolutionary time scales—rapid satDNA divergence is associated with genetic incompatibilities between closely related species (e.g.^9,10^). Even within species, individuals can vary widely in their abundance of satDNAs^11-13^. It is unclear what drives these evolutionary dynamics. Mutational properties of tandem repeats differ from single copy DNA: unequal crossing over, replication errors, and gene conversion can lead to rapid changes in sequence and abundance over short time periods^14-16^. SatDNAs can get caught in genetic conflicts over transmission to the next generation. Some satellites have the capacity to proliferate in genomes and bias their inheritance through selfish means^17,18^. For example, centromeric satDNAs can take advantage of the asymmetry in female meiosis and bias their transmission to the oocyte rather than the polar bodies, a process known as ‘meiotic drive’^19^. The tendency toward meiotic drive may explain why centromeric sequences, and the proteins that interact with them, are essential for chromosome segregation during cell division, yet evolve rapidly ^20^. SatDNAs can also become *targets* of selfish genetic elements like meiotic drivers in the male germline^5,21^.

Experimental model systems can help us explore the connection between satellite sequence or structure and functional effects. Functional assays of satDNAs across species have revealed phenotypic effects on embryo development^9^, nuclear organization^8^, sperm chromatin, heterochromatin formation^22^ and chromosome pairing^23^. In most cases, studying these phenotypic effects involve large chromosomal deletions^24-26^, chromosomal rearrangements^23^, or natural variation, each with different genetic backgrounds that make it difficult to attribute any effects specifically to the satDNA. Gene editing technology may help address this caveat: CRISPR-based approaches are effective at targeting sites in repetitive genome regions for selective chromosome elimination (*e*.*g*., Y chromosome in human cell lines ^27^ or to make large deletions of heterochromatin (*e*.*g*., Y deletions in Drosophila ^28^). We need ways to precisely interrogate these highly repetitive genome regions at the locus level.

Here we use the *Responder* (*Rsp*) satellite DNA of *Drosophila melanogaster* as an experimental model for understanding properties of pericentromeric satDNAs and their functional effects. *Rsp* is the target^29,30^ of a selfish male meiotic drive system called *Segregation Distorter* (*SD*)^31^. *SD* is a sperm-killer that is widespread in natural populations of *D. melanogaster* across the globe—it can spread in populations despite generally being costly for its bearer (reviewed in ^5^). *SD* chromosomes spread by achieving biased transmission through the germlines of heterozygous *SD/+* males by inducing a chromatin condensation defect in wild type sperm through an unknown mechanism^5^. The *SD* sperm-killing phenotype is positively associated with *Rsp* copy number: large alleles with many *Rsp* copies are highly sensitive to drive and small alleles with few copies escape drive^29,30^. We do not yet understand why *Rsp* is a target of drive. *Rsp* has no known molecular functions and is not essential^26^. Most alleles of *Rsp* in natural populations are large and therefore presumably sensitive to *SD* drive^11^. It is unclear why *Rsp* persists in populations when *SD* is so widespread. Previous work suggests that *Rsp* copy number is positively associated with fitness^32^, however this was based on a large X-ray-mediated deletion that removed a large block of pericentric heterochromatin^26^. To attribute the functional effects on spermatogenesis or fitness specifically to the satellite, we need targeted approaches based on a precise understanding of satDNA organization.

It has been challenging to understand satDNA organization—sequencing and assembling highly repetitive genome regions has only been possible recently^33^ with the advent of long read genome sequencing. We now know much about the detailed structural organization of the *Rsp* locus based on long-read sequence assemblies^34^. The *Rsp* satellite array primarily consists of tandemly repeated dimers of two related 120-bp repeats in the pericentromeric chromatin of chromosome 2R^29,30^. Within the *Rsp* locus, the tandem repeats are interspersed with transposable elements (TEs). The *Rsp* locus is flanked by simple tandem satellite repeats (AAGAG) on the centromere proximal side and TEs on the distal side^34^. Resolving the sequence and structural organization paves the way for designing targeted mutation experiments.

Here we present a CRISPR/Cas9-based strategy to generate an allelic series of mutations (deletions/expansions) of satDNAs using *Rsp* as our target repeat. Our approach involves a sequential CRISPR assay that targets complementary sites distributed throughout the locus. We demonstrate that this approach is highly efficient whether using an injection or transgenic strategy and in different genetic backgrounds of laboratory and naturally-derived strains. We use our mutation series to demonstrate the impact of *Rsp* structural mutations on the sperm-killing drive phenotype. Our study opens the door to precise manipulation of a compartment of the genome associated with critical phenotypes but inaccessible to traditionally genetic approaches.

## Results

### CRISPR-mediated manipulation of a satellite DNA locus

We focus here on the *Rsp* locus because of its functional relevance during spermatogenesis associated with copy number ^29,30^ and presumed effects on fitness^32^. Our earlier work resolved the structure of the *Rsp* satellite DNA array in the *Iso-1* reference strain using long reads, although this assembly is missing an estimated ∼22 kb of the proximal *Rsp* locus^34^. Here, we sequenced and assembled the *Rsp* locus in our *Cas9; Iso1* genetic background to serve as a basis for comparison. We confirmed the organization of the *Rsp* locus. *Rsp* is has a higher-order repeat structure consisting of tandem dimers of the left and right *Rsp* repeats, with interspersed transposable element (TE) islands (Figure 1A). Our assembly of *Cas9; Iso1* fills the gaps in previous assemblies and recovers additional sequence. *Rsp* has a higher-order repeat structure consisting of tandem dimers of the left and right *Rsp* repeats, with interspersed transposable element (TE) islands (Figure 1A). Our assembly of *Cas9; Iso1* fills the gaps in previous assemblies^34,35^. This reference allele is sensitive to *SD*. We designed guide RNAs based on this detailed organization, selecting guides that target complementary segments of the *Rsp* array with limited putative off target predictions (Figure S1; Table S1).

**Figure 1.**
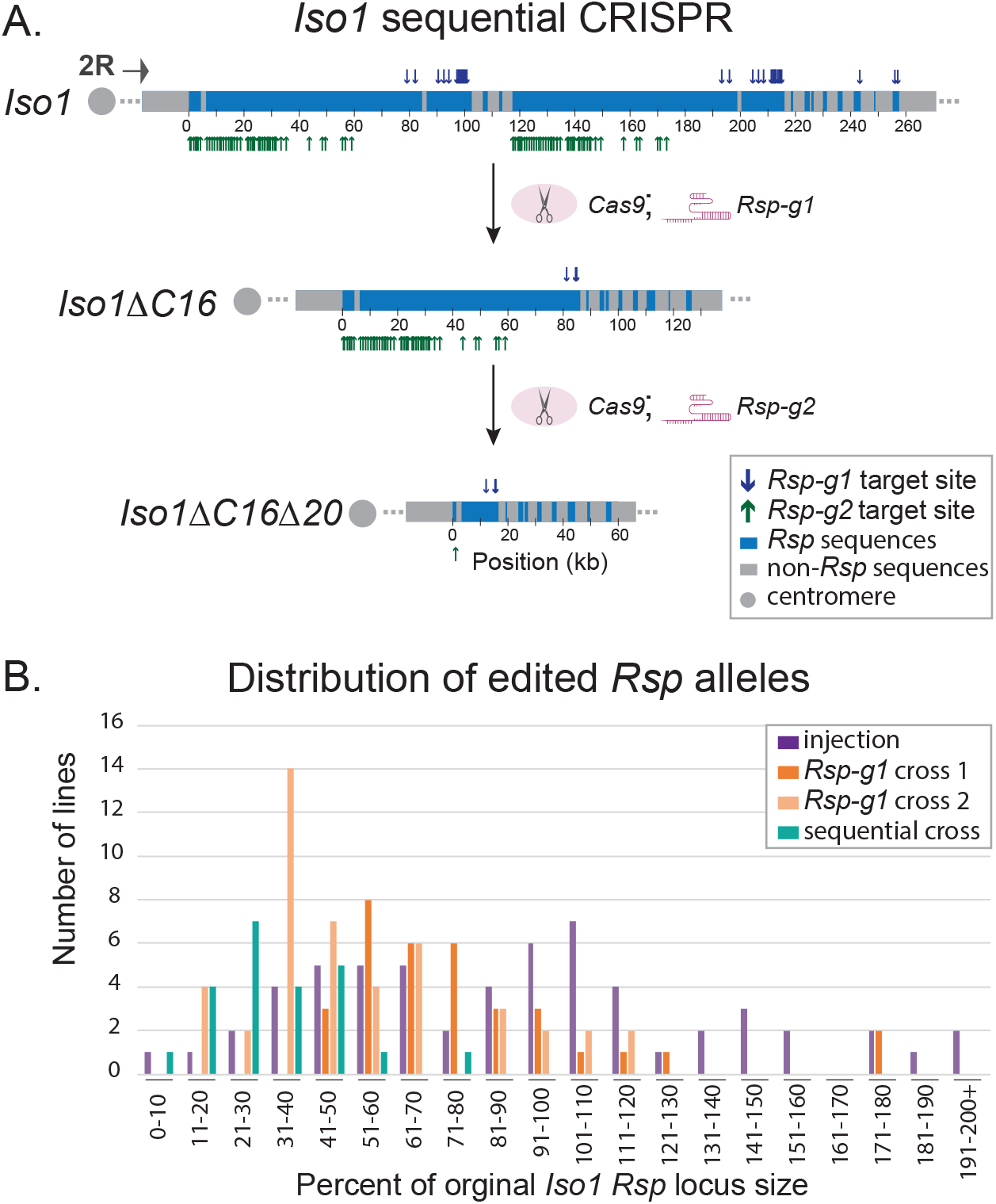
A sequential CRISPR scheme for targeted editing of the *Rsp* satellite locus. A) We implemented a CRISPR scheme to make sequential edits to repetitive loci and demonstrate its utility with the *Rsp* satellite—a complex satDNA in the pericentric heterochromatin of 2R in *D. melanogaster*. The scheme involves a set of crosses (Figs S2, S4) to establish stable fly lines with altered *Rsp* loci (panel B) and no changes in flanking sequences. We indicate target site locations for two guide RNAs with complementary distributions along the *Rsp* locus—*Rsp-g1* (blue down arrows) and *Rsp-g2* (green up arrows)—of the *Iso1* parent allele, a subline with a putative deletion (*Iso1*Δ*C16*) following CRISPR with *Rsp-g1*, and a subline with a large deletion (*Iso1*Δ*C16*Δ*20*) following sequential CRISPR with *Rsp-g2*. Both gRNAs target sites specifically matching *Rsp* repeat variants (blue) and avoid non-*Rsp* sequences in the locus (grey). Editing with this scheme occurs at high efficiency: few guide target sites remain following CRISPR (3/31 *Rsp-g1* in *Iso1*Δ*C16;* 1/99 *Rsp-g2* in *Iso1*Δ*C16*Δ*20*). B. We obtain a similar distribution of *Rsp* allele sizes after the first round of CRISPR editing, regardless of the method— injection or crossing (with two independent *Rsp-g1* experiments—cross 1 and cross 2; see Fig S2). The sequential CRISPR cross shifted the resulting allele distribution towards smaller sizes.

We implemented two complementary approaches to make *Rsp* mutations. Our first approach is based on injecting a guide RNA-expressing plasmid (*Rsp-g1*) targeting *Rsp* repeats, into embryos of the transgenic *Cas9; Iso1* strain (*Cas9; cn bw sp; +*; see methods and supplementary material). Following embryo injection, we implemented a crossing scheme (Figure S2) to obtain homozygous stocks with potential CRISPR-mediated mutations of the *Rsp* satellite locus. Six injected embryos produced fertile progeny that we used to create 59 homozygous stocks. We detected any differences in *Rsp* copy number with quantitative PCR (qPCR).

Our second approach involved crossing the same transgenic *Cas 9; Iso1* strain with transgenic flies expressing the *Rsp* guide RNA (*v1/Y; +/+; Rsp-g1/Sb*; see methods). We captured potential *Rsp* mutations over balancer chromosomes and created a total of 34 homozygous stocks containing potential mutations in the *Rsp* locus (Figure S2). We repeated the same crosses and generated 46 additional homozygous 2^nd^ chromosome lines. We screened these 80 stocks for *Rsp* copy number differences using a semi-quantitative PCR (*i*.*e*., comparing relative signals after a minimal number of cycles).

The different approaches and replicate experiments produced similar distributions of *Rsp* copy number across the 139 homozygous 2^nd^ chromosome lines relative to the original *Cas9; Iso1 Rsp* locus (Figure 1B). This suggests that the different CRISPR-based strategies produce a similar spectrum of *Rsp* mutations. We noticed an interesting pattern in the resulting alleles from the two different schemes: 1) we did not get a uniform allelic series of mutations, instead we tended to get discrete types of events; and while the majority of these single-guide CRISPR generated stocks were deletions, we did recover expansions with each scheme.

We selected 21 of the 139 lines for additional *Rsp* copy number estimation with semi-quantitative PCRs and quantitative slot blot analysis (Table S2). We selected one subline, *IsoΔC16* with an ∼50% deletion, for a preliminary assessment of CRISPR efficiency by sequencing with PacBio Hifi. Of the 31 potential *Rsp-g1* target sites in the *Cas9; Iso1* strain, only 3 target sites remained in *IsoΔC16*, suggesting that the CRISPR targeting was highly efficient. These results also suggest that additional crosses with *Rsp-g1* were unlikely to generate larger deletions. We, therefore, designed new guide RNAs targeting different sequence variants within the *Rsp* array and increasing numbers of target sites (*Rsp-g2*, 99 sites; *Rsp-g3*, 121 sites; and *Rsp-g4*, 506 sites) (Figure S1). Our initial crosses with each of these three *Rsp* guides with the *Cas 9; Iso1* strain resulted in few/no F1 progeny (see supplementary materials; Figure S3). Our subsequent crosses were consistent with the assumption that too many CRISPR-targeted cleavages can overwhelm DNA repair ^28^, but lethal off-target effects cannot be eliminated.

To create larger deletions of the *Rsp* array we, therefore, designed a sequential cross scheme that: *1)* takes advantage of balancer chromosomes to limit the possibility for recombination; and *2)* targets additional—but a moderate number of—sites (Figures S4; Figure 1).

Following the sequential CRISPR screen with a second guide RNA to target new sites within the *Iso1*Δ*C16 Rsp* locus (Figure S4), we generated 19 homozygous viable strains with putative sequential deletions (Fig S5; Fig 1B). We identified one subline, *Iso1C16*Δ*20*, with few *Rsp* repeats by molecular characterization (qPCR; Fig S5). We selected the following sublines as our allelic deletion/expansion panel for further genomic and functional analysis based on their *Rsp* alleles: *Iso1*Δ*C8 (deletion from injection), Iso1*Δ*C16* (deletion from single-guide cross), *Iso1ExpC11 (expansion from single-guide cross)*, and *Iso1*Δ*C16*Δ*20* (deletion from sequential CRISPR).

To demonstrate that our approach is useful beyond lab strains, we used our sequential CRISPR scheme (Figure S4) in an independent genetic background derived from a population in North Carolina^36^: *Ral309*. We generated an ∼20% deletion we refer to as *RalΔM17b* (Figure S6). To determine the structural organization of the allelic deletions/expansion, we used Oxford nanopore sequencing on each line and subline in our panel.

### CRISPR-mediated structural variations in the Rsp satellite: large deletions, expansions, and structural rearrangements

To determine the detailed structure of the allelic *Rsp* mutations, we *de novo* assembled each of our focal sublines and their respective parent strain used in the CRISPR screen. We have a contiguous and complete *Rsp* locus for each *Iso-1-*derived subline: the centromere proximal side of the *Rsp* array terminates in ‘AAGAG’ tandem satellite repeats and the distal side of the array terminates in TEs flanked by the tandem *Bari-1* repeat array ^34^(Figure S7).

Our assemblies of each *Iso1*-derived subline confirm *Rsp* deletions/expansion in an *Iso-1* genetic background with a high efficiency of targeted guide sites cleaved (Figure 1). They also confirm that we precisely and specifically, yet differentially, targeted the *Rsp* su*bline* loci while leaving all surrounding distal and proximal regions intact. In our parent *Cas9; Iso1* strain, there are two small islands of *Jockey*-type non-LTR retroelements (*G5*) interspersed with *Rsp* repeats, the array center has homogenized *Rsp* repeats consisting of an imperfect dimer of 120-bp left and right *Rsp* repeats, and the array edges have *Rsp* sequence variants, truncations, and TE insertions (Figure S7). Our sequencing confirmed two deletions of similar size (*Iso1ΔC8* ∼51.5% deletion by CRISPR injection and *Iso1ΔC16* ∼53.8% deletion by CRISPR cross scheme 1), one large deletion by sequential CRISPR cross scheme 2 (*Iso1Δ*C16*Δ*20 ∼88% deletion), and one expansion (*Iso1*ExpC11 ∼80.5% expansion by CRISPR cross scheme 1) (Figure S7; Figure 2).

**Figure 2:**
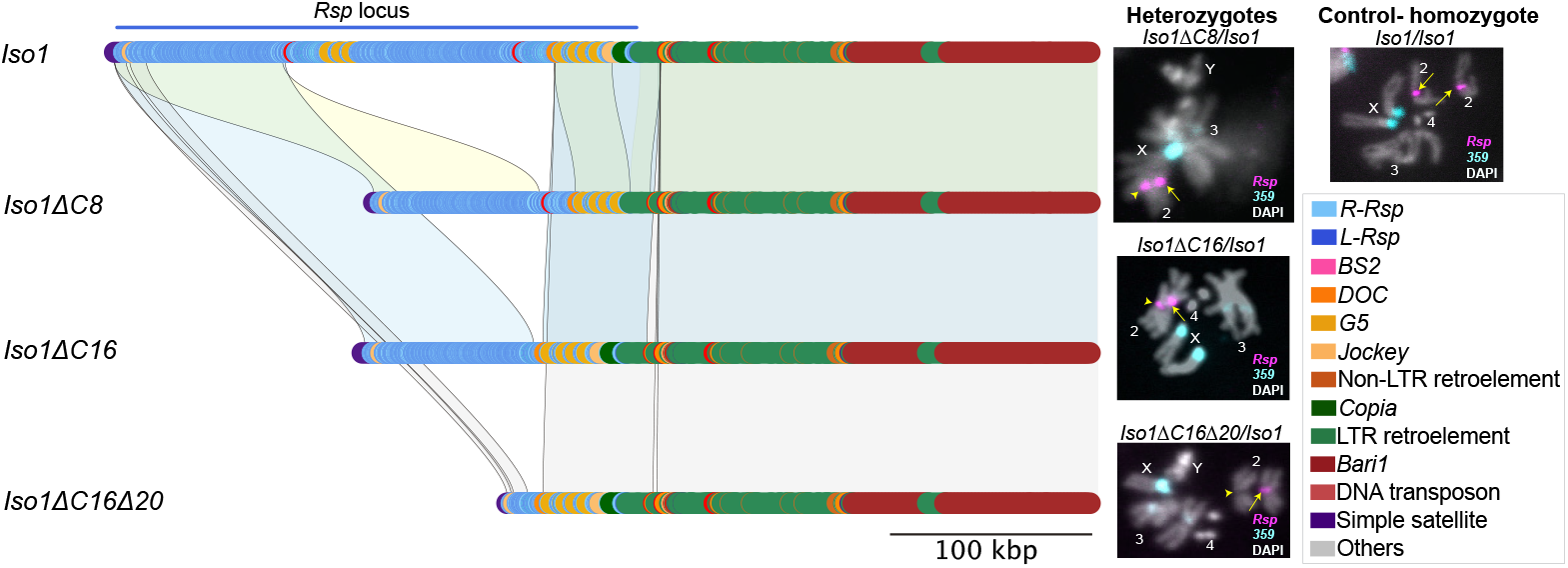
Synteny between *Iso1* and its sublines reveals simple deletions. The colored karyoplot shows the detailed structure of the genomic region containing the *Rsp* locus in the parent strain *Iso1* (total *Rsp* copy number 1756), and its allelic deletion series: *Iso1ΔC8* (total *Rsp* copy number 851), *Iso1ΔC16* (total *Rsp* copy number 812), and *Iso1ΔC16Δ20* (total *Rsp* copy number 211). In each subline, *Rsp* is flanked proximally by the simple satellite AAGAG and distally by TE clusters, most notably, the *Bari1* cluster. The shaded blocks between subline alleles indicate regions of pairwise synteny with *Iso1*: yellow blocks for *Iso1* and *Iso1ΔC8*; blue blocks for *Iso1* and *Iso1ΔC16*, and grey blocks for *Iso1* and *Iso1ΔC16Δ20*. In each case, the deletions appear to be simple: the distal *Rsp* repeats are deleted in both *Iso1ΔC8* and *Iso1ΔC16* and the *Iso1ΔC16Δ20* subline has further deletions of proximal *Rsp* repeats. The regions directly flanking the tandem *Rsp* repeats are largely unaltered in each subline. Fluorescence in situ hybridization (FISH) with *Rsp* probes and a control satellite not targeted by CRISPR (*359-bp*) in flies heterozygous for *Iso1* and the deletion subline allele, confirm that the *Rsp* foci are demonstrably smaller in *Iso1ΔC16* and *Iso1ΔC16Δ20*, consistent with their deletion sizes. The *Iso1/Iso1* homozygote is the control. Arrows point to the *Iso1 Rsp* allele and arrowheads to the subline *Rsp* allele.

*Iso1ΔC8* and *Iso1ΔC16* are nearly identical deletions of a distal island of *Rsp* repeats within the major *Rsp* locus, and a near complete elimination of target *Rsp*-g1 sites (Figure 1), a striking result given the different methods used to generate these deletions. *Iso1*Δ*C16Δ20* has an additional deletion of a proximal island of *Rsp* repeats, leaving only distal *Rsp* repeats interspersed with *G5* intact (Figure S7).

Our analysis of synteny suggests that the large deletions are simple in nature, corresponding to the presence of guide sites, without additional rearrangements (Figure 2). The origins of the *IsoExpC11* allele are interesting: these flies resulted from CRISPR in fly heterozygous for *Iso1*. After close inspection of synteny between expanded *IsoExpC11* and *Iso1* strain, it appears that *Iso1ExpC11* results from targeted recombination at the *Rsp* locus between two parent alleles. Although it is difficult to unambiguously resolve the events that led to this organization (Fig S8), we hypothesize that during the repair of the CRISPR-mediated double stranded breaks, the *Iso1 Rsp* locus recombined with the 2^nd^ chromosome in the *Rsp-g1* strain, in which nearly the entire 2L chromosome arm recombined (Table S3). Regardless of the exact nature of the expansion, we confirmed that regions distal to *Rsp* major locus on 2R have perfect identity and synteny to *Iso1* (Figs S7-S8, Table S3), suggesting that this was a targeted recombination event at *Rsp*. We confirmed structural aspects of the locus using PCR for junctions between *Rsp* and *G5* and *BS2* elements (see supplemental materials) and relative copy number by fluorescence in situ hybridization (Figure 2).

We sequenced both *Ral309* and *RalΔM17b* (∼20.5% deletion) and inferred the organization of each *Rsp* locus, although the assemblies are not contiguous or complete. We detect large structural variation in the organization of the *Rsp* major locus between *Iso1* and *Ral309*. Similar to *Iso1*, the *Ral309* major *Rsp* locus is flanked by tandem AAGAG satellites and the Bari-1 tandem array, however these alleles differ in their arrangement of interspersed TEs and the organization of *Rsp* variants. For example, *Ral309 Rsp* does not have the proximal G5 cluster, but does have insertions of other Jockey-type non-LTR retroelements like *BS2*. Based on synteny with *Ral309, RalΔM17b* organization suggests that the primary deletion occurred in the array center (Fig S9). As with the *Iso1*-background deletions, the mutations are limited to the *Rsp* locus—the sequences flanking the *Rsp* array are identical between the parent and deletion strain (Fig S9). Note that we did not produce a large deletion in *Ral309* like we did in *Iso1*. The sequence and structural divergence between *Ral309* and *Iso1* alleles lead to differences in the distribution of guide RNA target sites, with the crucial difference most likely being the presence of two distinct clusters of guide 1 target sites in Iso1 (Figure S6C).

Our sequencing allows us to identify putative off target effects of CRISPR on the 2^nd^ chromosome. Out of 10 predicted putative off target sites on chromosome 2, none show evidence for mutations in any sublines (both for *Iso1* and *Ral309*), suggesting that off target mutations are generally limited. We did detect several differences between these strains outside of the *Rsp* locus that are unrelated to our CRISPR scheme: *1)* in sublines originating from heterozygotes without a balancer chromosome to suppress recombination, we can detect recombination events on chromosome *2L* unrelated to CRISPR/Cas9 activity (*e*.*g. Iso1ΔC16* and *IsoExpC11*; Table S3); and *2)* we detected some variants between sublines that presumably accumulated since the lines were created (Table S3). None of the predicted variants outside of the recombination windows occur in coding regions of genes.

Taken together, our results demonstrate that the sequential CRISPR-based strategy targets complex satDNA arrays for deletion and expansion precisely and efficiently in both laboratory and wild-derived strains of *D. melanogaster*. Our approach therefore allows for the controlled manipulation of highly repetitive genome regions that have historically been refractory to genetic and genomic analysis.

### Consequences of CRISPR-mediation mutations for sensitivity to meiotic drive

Variations in satDNA can have functional impacts on chromosome behavior and organismal phenotypes^8,9,22,23,37^. *Rsp* is famous for being the target of the selfish male meiotic drive system, *Segregation Distorter* (*SD*)^29,30^, where sperm viability in males heterozygous for *SD* and a wild type chromosome depends on *Rsp* copy number^37^. Large alleles with many *Rsp* copies should be super sensitive to drive whereas small alleles with low copy number should be less sensitive to drive^29,30^. We therefore tested our alleles for drive sensitivity to determine if our CRISPR-mediated mutations have effects on spermatogenesis phenotypes.

While *Rsp* copy number should positively correlate with sensitivity to drive, many other loci across the genome can contribute to this phenotype, including the specific haplotype of the driver^5^. We therefore assayed the sensitivity of our *Rsp* allelic mutation series to *SD* drive in multiple genetic backgrounds and with multiple drive haplotypes. We homogenized the genetic background of each strain and controlled for *SD* chromosome effects on viability with reciprocal crosses. Some drive haplotypes can achieve perfect drive, where 100% of progeny from an *SD/+* heterozygous father inherit *SD* (*k* = 1.0) even with an “intermediate” *Rsp* copy number (Figure 3A; *SD5*). *Iso1* and the *IsoExpC11* subline with the expansion are both nearly completely sensitive to the strong meiotic driver *SD5* (Figure 3A). However, the deletion alleles—*Iso1ΔC16, Iso1ΔC8*, and *Iso1ΔC16Δ20*— each show reduced drive sensitivity (Figure 3A). To better differentiate drive sensitivity among *Rsp* alleles, we assessed differences in drive sensitivity with a weaker isolate of the *SD-Mad*^*1*^ haplotype. The sensitivity among sublines varied according to *Rsp* copy number, with the expansion allele being more sensitive to drive and the deletion alleles being less sensitive to drive (Figure 3B). We also explored the effects of the CRISPR mutations with an *SD-Mad* isolate of intermediate drive strength (Fig S10). We achieved similar results regardless of background except for *Iso1ExpC11* in *SD-Mad*^*2*^, where drive sensitivity is weaker than *Iso-1* despite the expanded *Rsp* allele. We may have introduced suppressor activity when swapping genetic backgrounds. *Iso1ΔC16Δ20* is interesting, as it demonstrates that alleles with only ∼200 repeats maintain some sensitivity (*k* > 0.5) to strong drivers (*SD5*, FET P=10^-8^), but not weaker drivers (*SD-Mad1*, FET P=0.18; *SD-Mad2*, FET P=0.11). Because *Ral309* and *RalΔM17b* are wildtype strains that do not have phenotypic markers to use in drive assays, we introduced a fluorescent marker to an *SD-Mad* chromosome and repeated the crosses. As expected, the *RalΔM17b* subline has lower sensitivity to drive (Fig 3C). Therefore, our CRISPR-mediated mutations that modify the structure and copy number of the *Rsp* locus result in phenotypic effects during spermatogenesis.

**Figure 3.**
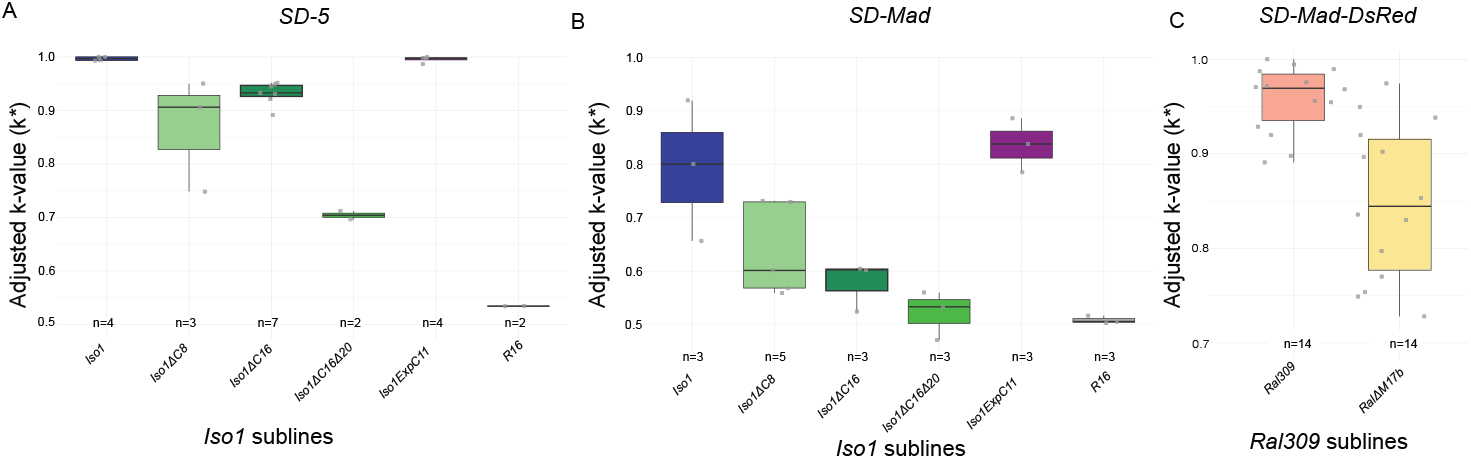
Edited *Rsp* alleles show altered sensitivity to meiotic drive. We tested each subline from the *Iso1* (A,B) and *Ral309* (C) allelic series for sensitivity to drive with drivers of different strengths and background. Drive strength, or *k*, is the proportion of *SD* offspring from heterozygous *SD/target* male, corrected for viability (by controlling for transmission through a heterozygous *SD/target* female). *R16* is a control—this line has a large deletion of pericentric heterochromatin including the *Rsp* locus and is completely insensitive to drive^26^. A. *SD-5* is a strong driver (*k*^*Iso1*^=0.996). The deletion alleles (*Iso1ΔC16, Iso1ΔC8, and Iso1ΔC16ΔC20)* each show reduced sensitivity to drive with *SD5*, while R16 is completely insensitive (FET P=0.41). The expansion *Iso1ExpC11* allele, like *Iso1*, remains highly sensitive to this strong driver. B. *SD-Mad*^*1*^ is an isolate of a weaker drive haplotype (*k*^*Iso1*^=0.792) that allows us to distinguish between the sensitivity at large copy number alleles. In this background, the deletion alleles (*Iso1ΔC16, Iso1ΔC8, and Iso1ΔC16Δ20)* show very little sensitivity to drive and R16 is completely insensitive (FET P=0.26), whereas the sensitivity of *Iso1ExpC11* is similar to *Iso1*, but shifted. Drive sensitivity with an intermediate-strength driver and background is in Fig S10. C. We repeated these crosses with a marked version of *SD-Mad* (*SD-Mad-DsRed*) and *Ral309* and its subline *RalΔM17b* and see reduced sensitivity to drive in the deletion allele. Taken together, our results indicate that CRISPR editing creates alleles with phenotypic effects on sperm viability.

## Discussion

Satellite DNAs are implicated in important cellular processes across taxa, but whether these effects depend on sequence composition, copy number, higher-order organization, or genomic context remain poorly understood. The highly repetitive nature and heterochromatic packaging^3^ of satDNAs, make them difficult to study at high resolution^33^. As a consequence, most functional studies of satDNAs involve large chromosomal aberrations or natural variation that confound the specific effects of satellite DNA with the surrounding genomic regions.

Disentangling the effect of a specific repeat from its genomic context has therefore been a key challenge. Here we overcome this limitation by using a sequential CRISPR strategy for targeted mutagenesis of specific satDNAs while leaving the flanking regions intact. We demonstrated that this approach is both precise and efficient when applied to the *Rsp* satellite in *D. melanogaster*: few target sites remain after editing and only the *Rsp* locus itself is altered. Notably, these targeted mutations have quantifiable phenotypic effects.

Our results have direct implications for the role of the *Rsp* satDNA in meiotic drive: we chose *Rsp* as our focal satellite because it is associated with a sperm chromatin phenotype and sensitivity to meiotic drive^29,30^. The evidence linking *Rsp* copy number to drive sensitivity has relied on the genetic analysis of recombinants and large structural mutations, and estimated differences in *Rsp* copy number among lab^29,30^ and natural strains^11^. While valuable, both data types are confounded by linked genetic variation and several suppressors of drive segregate in populations making it difficult to tease apart drive resistance at the *Rsp* locus from suppression. ^11^ By precisely manipulating *Rsp* in an otherwise intact background, we demonstrate that copy number changes at *Rsp* alone are sufficient to alter sensitivity to drive.

This work opens the door to functional analysis of satDNA and their molecular roles and contributions to fitness. A previous study from Wu and colleagues competed strains with deletions of the *Rsp* satellite against alleles with large copy numbers in population cages^32^. In the absence of *SD*, the large *Rsp* alleles had a fitness advantage over the deletions, suggesting that the *Rsp* satellite has a positive effect on fitness in the presence of *SD*^32^. A major limitation in that study, however, is that the deletion removed a large part of the heterochromatin on 2R^26^, so it is difficult to attribute any effects specifically to the *Rsp* locus. Our approach circumvents this limitation and provides a strategy for testing the functions and fitness effects of satDNAs like *Rsp* with locus-level precision.

This approach is best suited for repetitive loci with sequence-level information to aid in the design of target sites that fall into discrete, well-spaced, clusters. With the increasing availability of telomere-to-telomere genomes^38^, this approach can be readily extended to other repetitive loci. For example, targeting satDNA in essential genome regions like centromeres may help shed light on the role of DNA sequences in centromere function and the properties of repeats that contribute to these functions.

More broadly, our work establishes a framework to experimentally dissecting the roles of satDNAs at endogenous loci. Because the sequential CRISPR edits are made in the germline, allowing for the establishment of stable fly lines with the mutations, this provides a powerful approach to study satDNA contributions to chromosome dynamics, fertility, and fitness.

## Materials and Methods

### Selection of gRNA sequences and generation of gRNA expression vectors

We identified potential target sites and designed putative guide RNAs (gRNA) based on the major *Rsp* locus reported in Khost et al 2017^34^ using the CRISPR Optimal Target Finder tool (^39,40^; https://flycrispr.org/). We then mapped the recommended targets to the assembly and selected four guides with complementary target site patterns across the *Rsp* locus. We cloned each gRNA using expression vector pCFD3-dU6:3gRNA (Addgene plasmid #49410; ^41^) and the following primers: *Rsp-g1* sense: 5’-GTCGTACCCAAAAATAATTTGAATGG-3’ /antisense: 5’-AAACCCATTCAAATTATTTTTGGGTA-3’; *Rsp-g2* sense: 5’-GTCGTTTGTCTGGTTCTTGAAAT-3’/antisense: 5’-AAACATTTCAAGAACCAGACAAA-3’; *Rsp-g3* sense: 5’-GTCGAATTTCCGATTTCAAGTAAT-3’/antisense: 5’-AAACATTACTTGAAATCGGAAATT-3’; *Rsp-g4*: 5’-GTCGAGTTGAACAGAATCTCTAGA-3’/ antisense: 5’-AAACTCTAGAGATTCTGTTCAACT-3’. For *Rsp-g1* and *Rsp-g2*, we first 5’ phosphorylated primers with with T4 PNK (NEB). For Rsp-g3 and Rsp-g4, we phosphorylated primer pairs with Integrated DNA Technologies. We annealed paired phosphorylated sense/antisense primers at 37°C and then ligated into BbsI digested pCFD3 (50 ng) with T4DNA ligase. We transformed each ligated vector into competent bacteria (Trans1-T1Phange resistant chemically competent cells; www.transbionovo.com) and plated on LB Ampicillin plates. We extracted DNA from colonies using a plasmid DNA mini kit (OMEGA E.Z.N.A Plasmid DNA Mini Kit I) and verified each insert was verified by sequencing (ACGT, Inc.).

### Cas9 fly strain

The original *D. melanogaster Cas9* strain (RRID:BDSC51323) expresses the Cas9 protein in the ovary under control of *vas* regulatory sequences (*y[1] M{RFP[3xP3*.*PB] GFP[E*.*3xP3]=vas-Cas9}ZH-2A w[1118]*). We introduced the X chromosome from this *Cas9* strain into the sequenced strain, *Iso1* (Bloomington Stock 2057,^42^) to make flies homozygous for the *vas-Cas9* X chromosome as well as homozygous for *cn bw sp* (from *Iso1* 2^nd^ chromosome): *w y [vasCas9 GFP RFP]; cn bw sp; +* (hereafter *Cas9; Iso1*) see Figure S2, S4). We repeated this process to introduce *Cas9* into a wild-derived strain from Raleigh, NC, *Ral309* (RRID: BDSC 28166) to make *Cas9; Ral309*.

### Generating deletion/duplication lines via CRISPR

We injected the *Rsp-g1* vector (above) and pcFD3Ebony1 (Addgene #83380; ^43^) into our *Cas9; Iso1* strain using GenetiVision services (www.genetivision.com). We screen injected adults by individually crossing to *w*; *CyO/Gla*, and we monitored homozygous 2^nd^ chromosome sublines for changes in *Rsp* copy number (see Figure S2) by PCR, qPCR, and slot blots (see below).

We generated transgenic flies expressing each guide (*Rsp-g1* through *Rsp-g4*) from the PhiC31 docking site *VK05 [(3L) 75A10]* using GenetiVision services and balanced with *TM3Sb*. We maintained our guide RNA stocks as +;+/CyO;*Rsp-gRNA/TM3Sb*. For CRISPR, we crossed homozygous *Rsp-gRNA CyO* males to virgin females from our *Cas9; cn bw sp* stock or the *Cas9; Ral309* stock. We then individually cross the F1 *CyO* male progeny to virgin *CyO/Gla* females, and generated single ‘modified’ 2^nd^ chromosomes stocks (Figure S4). We screened each subline for changes in *Rsp* copy number.

### Monitoring for changes in Rsp copy number

We estimated the size of the *Rsp* locus in sublines generated in the CRISPR crosses with a combination of approaches. First, we did a semi-quantitative PCR assay with a minimal number of number of cycles, and examined the relative intensity to controls (*e*.*g*. the parent *Cas9; Iso1* or *Cas9; Ral309* line) and analyzed multiple sibling replicates. Briefly, we did single female ‘squish preps’^44^ in 50 ul buffer (10 mM Tris-Cl pH 8.2, 1 mM EDTA pH 8.0, 25 mM NaCl, 200 ug/ml proteinase K) at 42°C for 30 minutes, and then boiled for 1 minute^44^ and used a 10-fold dilution in a NEB standard PCR reaction with primers *Rsp1* 5′-GGAAAATCACCCATTTTGATCGC-3′ and *Rsp2* 5′-CCGAATTCAAGTACCAGAC-3′ and Taq polymerase for 15 cycles. We determined band intensities on 1% agarose gels on a ChemiDocXR+ using ImageQuant.

We selected sublines with consistent differences in *Rsp* signal from the controls with qPCR and/or slot blot. For qPCR, we surveyed individual females (in triplicate) using the squish protocol above and used a 20-fold dilution in a standard BioRad iTaq Universal SYBR Green reaction. We normalized *Rsp* (with same primers as above) to the *tRNA-Lys-CTT* (tRNA forward 5’-CTAGCTCAGTCGGTAGAGCATGA-3’ and tRNA reverse 5’-CCAACGTGGGGCTCGAAC-3’) to determine relative abundance.

For slot blotting, we mashed five female flies (in triplicate) in 200 ul of squish buffer. To extract nucleic acids, we used an equal volume of phenol/chloroform/isoamyl alcohol (1:1:0.01), vortexed for 45 - 60 seconds, and then centrifuged for 3-5 minutes. We added an equal volume of chloroform/isoamyl alcohol (1:0.01) to the aqueous top layers, vortexed for 30 seconds, then centrifuged for 1 minute and then repeated this step and determined the concentration of total nucleic acid by nanodrop. We denatured ∼200 ng of the nucleic acid (final concentration 0.25 M NaOH, 0.5 M NaCl) for 10 minutes at room temperature, transferred to a tube with an equal volume of ice-cold loading buffer (0.1X SSC, 0.125 M NaOH). We loaded samples as recommended for the 48-well BioDot SF microfiltration apparatus (Bio-Rad), washed with 200 μl of loading buffer. To make the biotinylated RNA probes, we gel extracted PCR amplicons (primers: *Rsp* 5’-TAATACGACTCACTATAGGGGAAAATCACCCATTTTGATCGC-3’ and 5’-CCGAATTCAAGTACCAGAC-3’; rp49 5’-GTAATACGACTCACTATAGGGCAGTAAACGCGGTTCTGCATG-3’ and 5’-CAGCATACAGGCCCAAGATC-3’) and transcribed using the Biotin RNA Labeling Mix (Roche) and T7 polymerase (Promega). We rinsed the nylon membrane (GeneScreen Plus) for 2 minutes with 2X SSC before being UV crosslinked (Stratalinker) and then hybridized with a biotinylated rp49 RNA probe in North2South hybridization solution (ThermoScientific) at 65°C overnight. We processed as recommended for the Chemiluminescent Nucleic Acid Detection Module (ThermoScientific) recorded the signal on a ChemiDoc XR+ (Bio-Rad). We stripped the membrane with a 100°C solution of 0.1X SSC/0.5% SDS (3 times for ∼20 minutes each) and re-hybrized with a *Rsp* probe (60°C overnight) and processed as with rp49, and quantitated signals using ImageLab software (BioRad).

### Validating Rsp-TE boundaries with PCR

We designed primer sets that specifically amplify sequences at the junction of TE insertions and *Rsp: G5* clusters or *BS2* insertions, with PCR. We used the following primer pairs for G5: G5(5,3) FOR 5’-TCGATGAAGCTAATTGCTGG-3’ and G5(3,5) REV 5’GTGGTATGCCTAATGGGAG-3’. To distinguish the *G5-3* and *G5-5* clusters^34^, we used a restriction digest based on the *AhdI* site specific to the *G5-5* element and ran on a gel. Two bands in the digested product (2.0kb and 1.5 kb) indicate the presence of both elements; a single band of 1.5 kb suggests the presence of the *G5-5* element only; a single band of 2.0 kb suggests the presence of the *G5-3* element only; and the lack of either of those bands suggests neither is present. We validated *BS2-Rsp* junctions using PCR (*BS2-Rsp* For: 5’-TATTCGTTTTACTTCCGTCGTATAAAGGAAAAAGGTTCGT and *BS2-Rsp* Rev: 5’-GCCGATGGTTGATTTGGGTTTT).

### Genome sequencing and assembly

We isolated genomic DNA from 20 mg of 5-7 day old female flies using the NEB Monarch spin gDNA extraction kit (T3010S) following a modified protocol^45^. We sequenced the gDNA using the custom eukaryotic genome sequencing workflow at plasmidsaurus with Oxford Nanopore Technologies (ONT) R10.4.1 chemistry and multiplexed sequencing on two full flow cells: one for *Cas9; Iso1* and its derived allelic mutations (*Iso1ExpC11; Iso1ΔC16; Iso1ΔC8; Iso1C16Δ20*), as well as *Ral309* and its allelic mutation line *RalΔM17b*. Plasmidsaurus did basecalling of POD5 files using Dorado (super accurate model, SUP) version 4.3.0 on an NVIDIA A100 80Gb PCIe GPU^46^. We assembled the raw FASTQ files using hifiasm (version 0.25.0) with –l0 --ONT parameters and without read filtering. We converted the primary contig gfa files into FASTA format using *awk ‘$1 == “S” {print”>“$2”\n”$3}’ <filename*.*gfa> > <filename*.*fasta>*, and annotated repeats using RepeatMasker (version 4.1.0) with a custom *D. melanogaster* repeat library^35^.

### k-value assays

We assayed sensitivity to *SD* by crossing males heterozygous for *SD* and a target 2^nd^ chromosome ([target]/*SD)* to *Iso1* females and measuring drive strength (*k*) as the proportion of *SD* offspring. *k* ranges from 0-1 where 0.5 is Mendelian segregation and 1.0 is perfect drive. To introduce different genetic backgrounds, we backcrossed flies with our target 2^nd^ chromosomes to either y; CyO/Gla or *SD-Mad* for three generations. We corrected each *k* value for viability differences by adjusting ratios based on reciprocal crosses between [target]/*SD* females and *Iso1* males.

### Variant Calling

We mapped our ONT sequence reads to its respective parent de novo assembly (*Cas9; Iso1* or *Cas9; Ral309*) with minimap2 (version 2.26) and *–ax map-ont* parameters and converted the output to sorted BAMFILES using samtools (version 0.21.0). We used these sorted bamfiles to call variants using Clair3^47^ with *r1041_e82_400bps_sup_v430_bacteria_finetuned model* and *--include_all_ctgs* options using 2 H100 Nvidia GPUs in EmpireAI HPC cluster^46^. We filtered VCF files generated from Clair3 for QUAL > 30 and Read depth between 25% and 75% quantile range using vcfR (version 1.15.0)^47^package in R. We annotated the filtered VCF files using snpEff (version 5.4) ^48^ based on a custom database generated from *Cas9; Iso1* and *Cas9; Ral309* assemblies and genome annotation GFF files generated using liftoff ^49^ from our previously published annotations of the *D. melanogaster* genome^35^. To infer the recombination breakpoints, we used vcfR (version 1.15.0)^47^. Briefly, we extracted the information for genotype (GT), allele depth (AD), and read depth (DP) from filtered vcf files for individual chromosome 2 contigs. We calculated the alternate allele depth ratio using sum(alternate-AD/(reference-AD+ alternate-AD), assigned alternate, reference, and heterozygous based on the AD ratio of >=0.9, <=0.1, >=0.3 and <=0.7, and the rest as uncertain. We then used the frequency of alternate allele depth in a window of 50kb across the contig to infer putative recombination breakpoints.

### Fluorescence In Situ Hybridization (FISH)

We performed FISH as in Larracuente and Ferree^50^. Briefly, we dissected larval brains were in 1× PBS, treated with a hypotonic solution (0.5% sodium citrate) and fixed in 1.8% paraformaldehyde, 45% acetic acid, and dehydrated in ethanol. We labeled the *Rsp* satellite using Stellaris probes (0.125µM) and the *1*.*688* satellite as a control using an oligo probe (5’ FAM-TTTTCCAAATTTCGGTCATCAAATAATCAT;^51^). We hybridized at 95°C for five minutes and then overnight at 30°C, washed three times with 0.1× SSC, and mounted slides with SlowFade Diamond Antifade mountant (ThermoFisher, cat # S36963). We visualized slides on a Leica DM5500 upright fluorescence microscope at 100× outfitted with a Hamamatsu Orca R2 CCD camera and analyzed images using Fiji (ImageJ) software.

### Synteny

To generate the synteny plots, we used the main *Rsp* contigs from each genome assembly, annotated with Repeatmasker (see above), and aligned the *Rsp* contigs from each parent strain and its variant subline using *nucmer* from Mummer4 with *--mum* option^52^. We generated coords files from the nucmer output files using *show-coords –b* option. We generated synteny plots using KaryoploteR ^53^ package in R. For *Iso1* allelic series, any blocks that were ≤2000 bp were filtered out before plotting and the proximal *G5* synteny block was removed manually using Adobe Illustrator. For *Ral309*, any block ≤10000 bp were filtered before plotting and any overlapping blocks with the larger blocks were removed manually.

For *Ral309*, we generated superscaffolds based on a preliminary analysis of synteny between *Ral309* and *RalΔM17b*. We placed the proximal *Ral309* contig based on its alignment with the putative proximal *RalΔM17b* contig and on the presence of *AAGAG* sequences adjacent to the *Rsp*^34^. We placed the distal contig based on the presence of *Bari1* elements. To count the total number of *Rsp* repeats (Fig S7), we used blastn with consensus *L-Rsp* and *R-Rsp* sequences, designated Rsp units into variant and truncated based on the sequence length (90bp), bitscore (0.7), and percentage identity (90%)^34^.

### Creating a fluorescently-marked SD line

We created a fluorescently-tagged *SD* chromosome by inserting an *Sd-RanGap-3xFLAG* transgene inserted downstream of the CG10195 on the 2nd chromosome from a revertant SD-Mad strain lacking *Sd-RanGAP* ^54^ using CRISPR/Cas9. We generated the gRNA expressing construct by inserting the gRNA sequence (GAATTAACAGACAGCGCAGG) into the BbsI sites of the pCFD3 vector following the protocol in Port et al. ^41^. To generate the repair construct, we amplified the homology arms around the CRISPR cut site from *SD-Mad* genomic DNA and inserted the left homology arm (For 5’-CGCTGAAGCAGGTGGAATTCCCAGTGACAATGTGACATGCCGTAA;Rev 5’-GTGTGCATATGTCCGCGGCCGCTAGGTACCAACGGGAAGGTCGACAAAGCCGCC CATAAA) into the pDsRed-attp (Addgene #51019) vector with EcoRI and NotI and the right homology arm (For 5’-AAGAGCTCTAGAAAAGATCTCCTGCGCTGTCTGTTAATTAGAAAATCGGT; Rev 5’-AAGAGCCTCGAGCTGCAGAAGGCCTAGGCGTTCAAAGCATCTGCATCGCTATACCT) with SpeI and AvrII. We inserted the linker-3xFLAG and *Sd-RanGap* 3’UTR (For 5’-ACGATGACAAGTAGCCGCGGTCGGGCACTCGTAAGGCCAGCGCGACTTAA;Rev 5’-CGATCGCAGGTGTGCATATGTCGTTTACATTCTAGAACAAATATAGGAAG) into this construct with NotI and NdeI, then amplified the *Sd-RanGap* promoter and gene from *SD-Mad* genomic DNA (For 5’-TCGACCTTCCCGTTGGTACCTTAAATTATTTATATTTTGAATACCTTTTA; Rev 5’-CACTCCTGAATCGCCCGATGCGGCCGCTTCCAAATCCTCCTCTTCCTGGTCCTGAA ACGG) and inserted it upstream in-frame with the linker with KpnI and NotI. A fly stock was generated with the X chromosome from the Bloomington Drosophila Stock Center #51323 (y[1] M{vas-Cas9}ZH-2A w[1118]) and the 2nd chromosome from the revertant *SD-Mad* strain, the gRNA and repair template constructs were injected into embryos, DsRed fluorescent-eyed progeny were selected, and balanced over CyO by GenetiVision.

## Acknowledgments

We thank Danielle Khost and Stephanie Hao for help with fly stocks, Adya Mohapatra and Olivia Hullihen for help with PCR, and Zeenath Unnisa for help with the *Rsp-g1* plasmid. We also thank the Larracuente lab for feedback and Nathaniel and Helen Wisch for their support. This work was supported by NIH GM119515 to AML. We gratefully acknowledge use of the research computing resources of the Empire AI Consortium, Inc, with support from the State of New York, the Simons Foundation, and the Secunda Family Foundation. We thank the Center for Integrated Research Computing for access to computational resources.

## Data availability statement

All code to reproduce figures and analysis is available on Github (https://github.com/LarracuenteLab/Rsp.CRISPR) and figshare (DOI forthcoming). All sequence data are deposited in NCBI (accession numbers forthcoming).

## Notes

### Competing Interest Statement

The authors have declared no competing interest.

