## Supplementary materials for "Targeted editing of pericentromeric satellite DNA alters sensitivity to meiotic drive"

**CRISPR/Cas9-based approaches to generate satDNA mutations**

We implemented two complementary approaches to make *Rsp* mutations. First, we injected embryos (*Cas9; cn bw sp; +)* with a guide RNA coding plasmid (*Rsp-g1*) which targets the *Rsp* satellite array. We mated emerging adults (Figure S2) and obtained homozygous stocks with putative CRISPR-mediated mutations of the *Rsp* satellite locus. These embryos produced 59 homozygous stocks which were screened for *Rsp* copy number differences by qPCR.

Our second approach involved crossing the same transgenic *Cas9; Iso1* strain with transgenic flies expressing *Rsp* guide RNA (*v1/Y; +/+; Rsp-g1/Sb*; Figure S2; see methods). Emerging F1 flies were mated to the 2$^{nd}$ chromosome balancer strain *CyO/Gla* and the scheme followed (Figure S4) generating a total of 34 homozygous stocks containing potential mutations in the *Rsp* locus. These stocks were screened for *Rsp* copy number differences by semi-quantitative PCR (i.e. comparing relative signals after a minimal number of cycles). Biological replicates were performed. This cross was repeated- the only difference being that CyO was introduced into the transgenic *Rsp-g1* fly line (*v1/Y; +/CyO; Rsp-g1/Sb)* to eliminate crossing over on the 2$^{nd}$ chromosome (Figure S4). Cross 2 produced 46 homozygous 2$^{nd}$ chromosome lines; each of which were also screened for *Rsp* copy number differences by semi-quantitative PCR.

The *Rsp* copy number estimates for each of the 139 (59+34+46) homozygous 2$^{nd}$ chromosome lines were averaged and grouped by PCR signal relative to that of the original *Cas 9; Iso1 Rsp* locus. The distribution of estimate sizes is shown in Figure 1B. In general, the results between the three experiments are similar. However, the cross 2 results shift towards slightly larger deletions than cross 1- although this is more likely due to PCR fluctuations than a difference in deletion sizes. Deletions and expansions were observed in both approaches. The same range of changes were also generated during our sequential CRISPR scheme (Figure 4) using one of our generated deletion strains.

We next selected 21 of the 139 lines to reassess their estimated mutation sizes. Of particular interest were the lines in which the first round of PCR suggested the largest deletions- i.e. under 30% of the *Rsp* locus remained. We also reassessed multiple lines which appeared to have retained approximately half the *Rsp* repeats (the most common CRISPR event recovered), and several lines which seemed to have undergone an expansion. Additional semi-quantitative PCRs and quantitative slot blot analysis were performed on these 21 lines (Table S2). Five of these strains were also tested for their sensitivity to *SD,* as smaller alleles should be less sensitive to drive, and large alleles, more sensitive to drive (Table S2). The smallest allele that we generated with a consistent *Rsp* copy number estimate and the corresponding expected *SD* drive k value (proportion of *SD* progeny from *SD/*+ males) was *Iso1∆C16* at ~50%.

To examine the efficiency of our CRISPR scheme, we selected *Iso1∆C16* for long read sequencing with PacBio Hifi. Comparison of the *Iso1∆C16* sequence assembly versus that of *Iso1*[1] revealed multiple junctions resulting from CRISPR target events. The largest deletion appears to remove sequences from the proximal *Rsp g-1* target site cluster to the beginning of the distal guide 1 target site cluster. over 100 kb away. An approximately 7 kb region with a cluster of 10 target sites has been pared down to an ~2.4 kb region in such a jumble that it is difficult to align what is left with the *Iso1* assembly. The three target sites which are significantly removed from the two clusters have also been cleaved. The first has 3 non-templated bp inserted at an otherwise clean junction. Therefore, out of 31 potential *Rsp-g1* target sites, only 3 target sites remain. While we obviously selected a strain that should have a large segment of the *Rsp* locus deleted, these results suggest that CRISPR targeting can be highly efficient.

The *Iso1ΔC16* sequencing results also indicated that additional crosses with *Rsp-g1* were unlikely to generate larger deletions. We, therefore, designed new guide RNAs targeting different sequence variants within the *Rsp* array with increasing numbers of target sites (*Rsp-g2*, 99 sites; *Rsp-g3*, 121 sites; and *Rsp-g4*, 506 sites) (Fig S1; Table S1). Our initial crosses with each of these three *Rsp* guides with the *Cas 9; Iso1* strain resulted in few/no F1 progeny (Figure S3). The lethality in the case of *Rsp-g2* was due at least in part to what we thought would be an infrequent off-target cut in the Myosin 28B1 gene. Crosses between strains with a decreasing number of *Rsp* repeats/target site with *Rsp-g2* or *Rsp-g3* gave rise to an increasing number of F1 progeny. However, crosses with *Rsp-g4* only gave rise to progeny when there were no *Rsp* repeats on either 2$^{nd}$ chromosome. We therefore designed a sequential cross scheme that takes advantage of balancer chromosomes to limit the possibility for recombination, and targets a moderate number of sites (Figure S4).

**Supplemental figures**

Fig S1

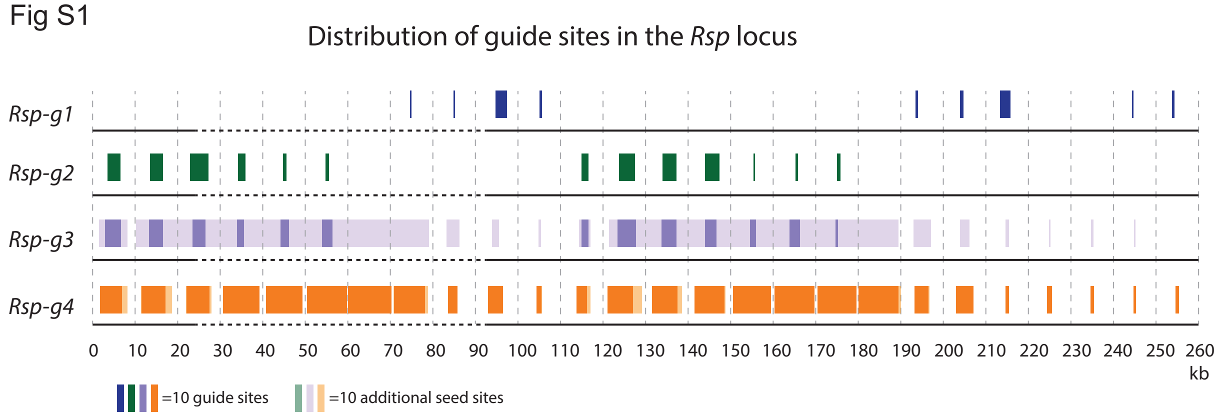


**Figure S1-Distribution of guide target sites**: The distributions of target sites across the *Rsp* array- in 10 kb intervals- of the 4 *Rsp* guides used are shown (dark boxes). Because off-target cleavages within the array at non-perfect sequences are possible, the distributions of seed target sites (light boxes) are also shown for the 3 guides (*Rsp-g2*, *Rsp-g3*, *Rsp-g4*) for which these additional sites occur.

### Fig S2

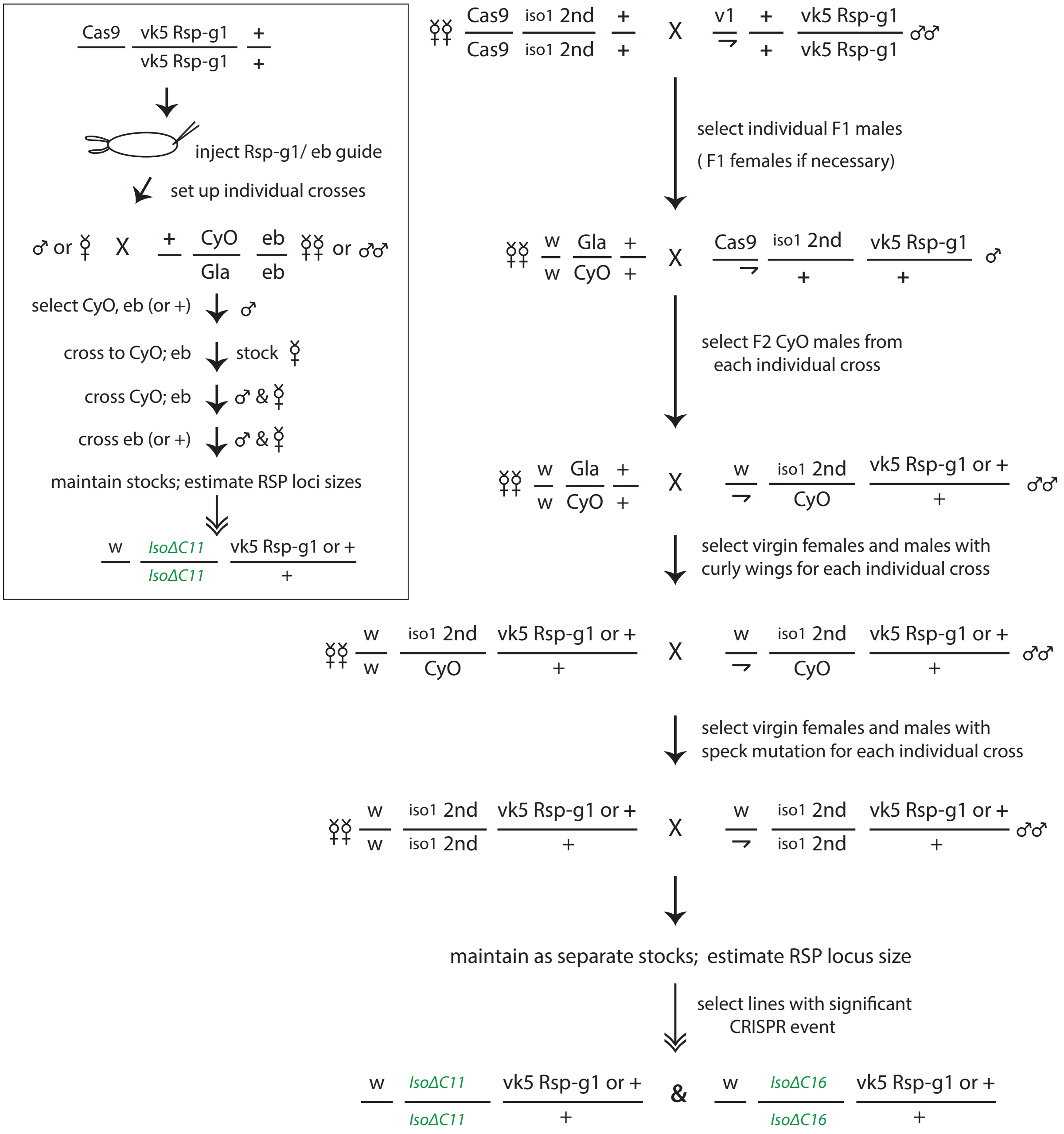


**Figure S2-Single-guide CRISPR cross scheme**: Originally *Rsp-g1* was introduced into our Cas9; *Iso1* line in alternative ways- by injection of a plasmid (summarized on the left) and by crossing to a second line with the guide RNA inserted on the third chromosome (summarized on the right). Adults emerging from the injected embryos were crossed to a balancer stock, and those progeny selected and crossed as diagrammed. F1 progeny arising from the cross on the right were selected and also mated to balanced flies as indicated. Flies which are *cn bw sp* (white eyes with a dark spot at the base of the wing) were selected because this implies homozygosity of chromosome arm 2R- the location of the *Rsp* array. Homozygous stocks derived from both injection and cross descendants potentially have a modified *Rsp*. Shown in green are 3 examples of strains coming out of these experiments.

Fig S3

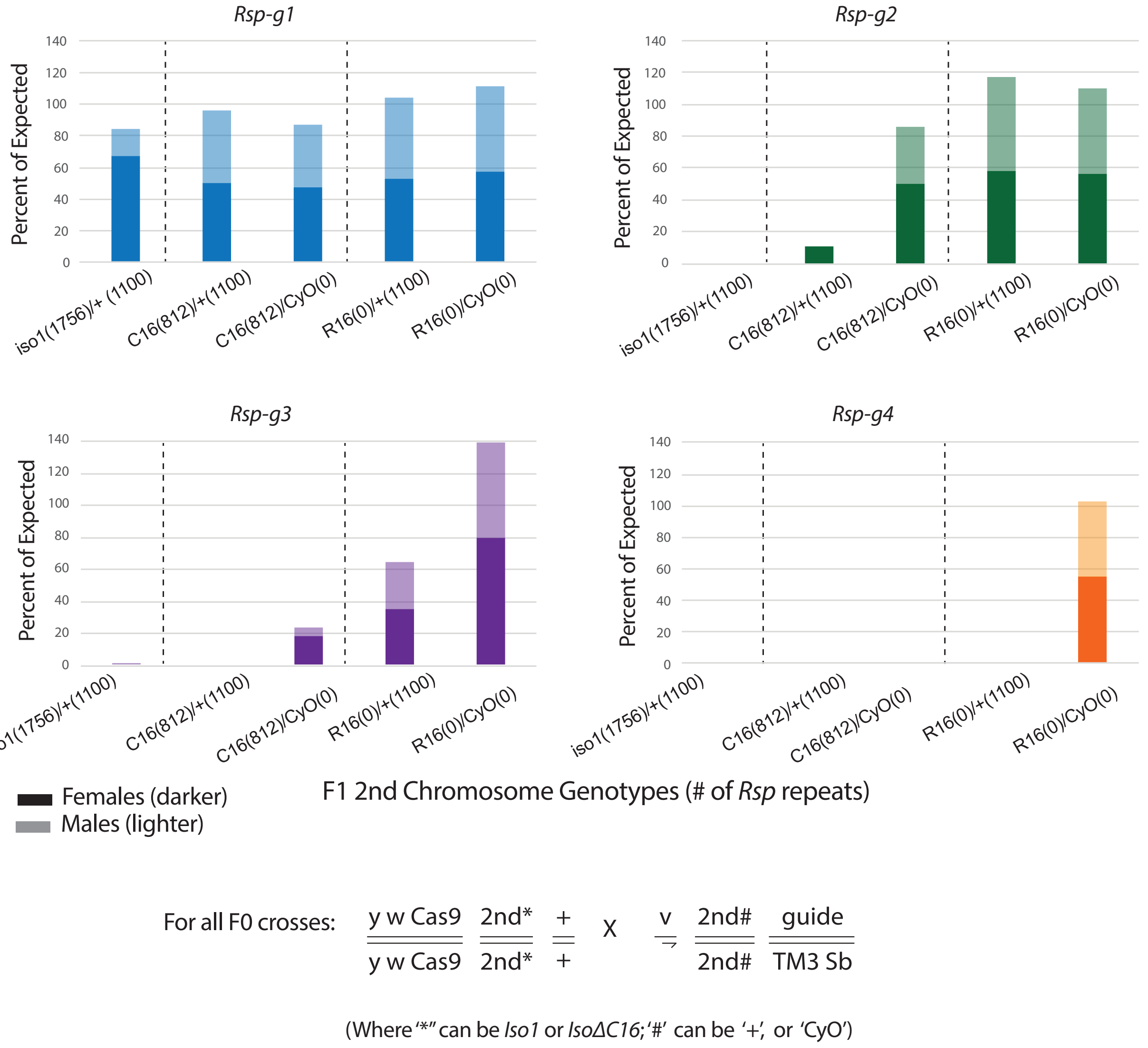


**Figure S3-Survivability in CRISPR crosses**: Because these crispr studies potentially involve cleaving many target sites within the *Rsp* array, it is not surprising that some progeny do not survive because the DNA repair mechanism is overwhelmed. To look at this, F1 progeny- males (light bars) and females (dark bars) separately- from the indicated crosses were counted and plotted above. Noted for each cross is the number of guide target sites on each of the second chromosomes. In general, fewer F1 males emerged. And across the crosses, it is also apparent that fewer F1 flies survive as the number of potential guide target sites increases.

Fig S4

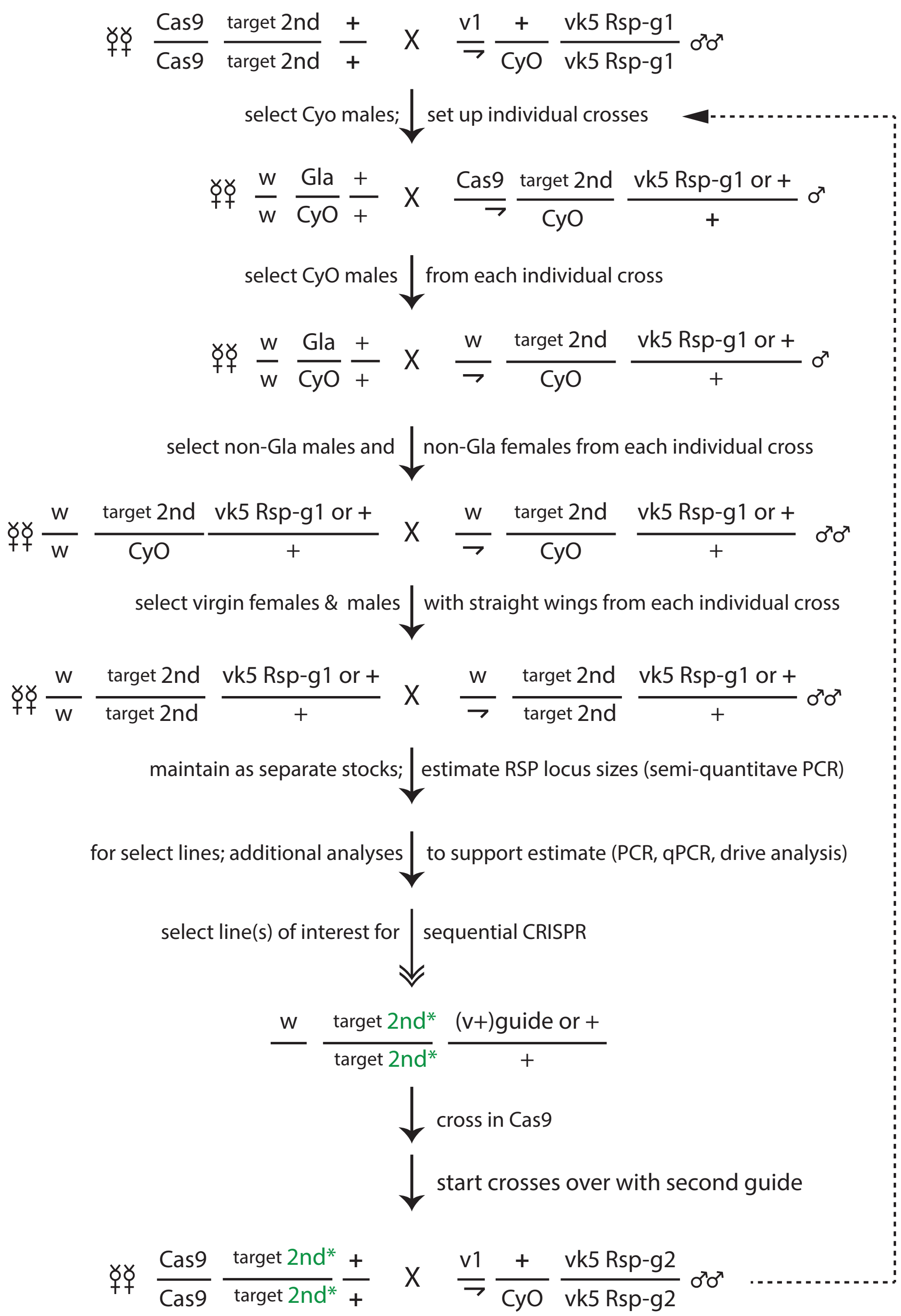


**Figure S4- Sequential CRISPR cross scheme**: The original crossing scheme (Figure S2) was tweaked after the first crosses so that the presence of the second chromosome balancer, *CyO*, would minimize crossing over during our CRISPR attempts. The sequential step generating *Iso1ΔC16Δ20* and the *Ral309* deletions were done using this scheme.

#### Fig S5

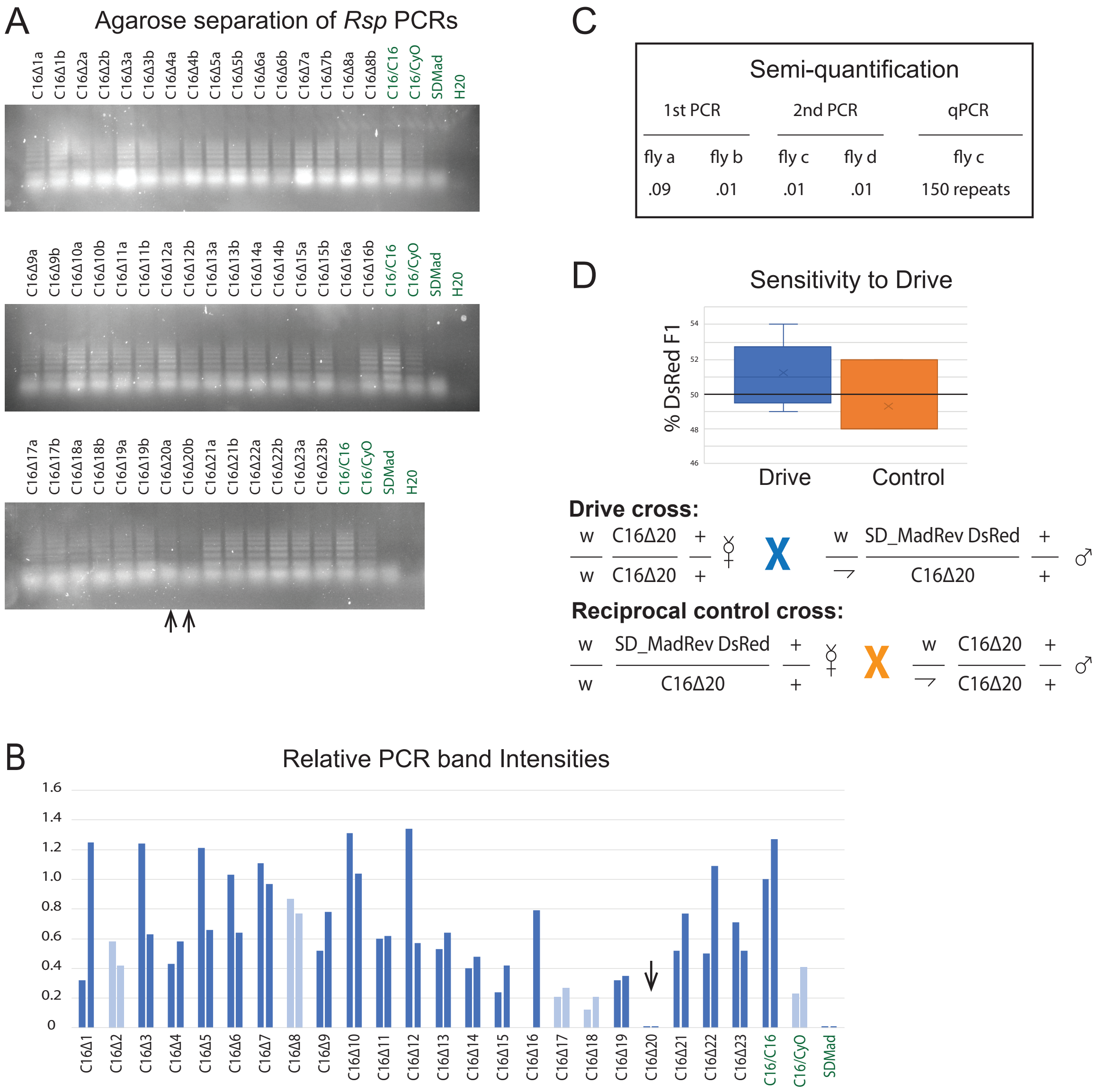


**Figure S5-Sequential CRISPR screening and Iso1∆*C16*∆*20***: A. The final step in *Iso1* sequential CRISPR (crossing *cas9;* I*so1∆C16* females to *Rsp-g2* males) generated 23 strains. Only 19/23 were fertile (dark blue boxes) when homozygous for the 2$^{nd}$ chromosome. All 23 strains were analyzed by our semi-quantitative PCR protocol, and the products separated on agarose gels. B. Using *Iso1∆C16* homozygous flies as standard, the relative intensities of the bands generated for each line were graphed. The *Iso1∆C16∆20* (arrows) comparisons suggested a relatively large deletion in the *Rsp* array of that strain. C. To support that conclusion, two additional flies were assessed in a second PCR. A qPCR (three technical replicates) for fly c was also done. PCR results are summarized as locus size relative to the *Iso1∆C16 Rsp* array size. The qPCR repeat number estimate is based on the 1756 repeats reported here. D. The projected small size of the *Iso1∆C16∆20 Rsp* array suggested it would not/only slightly be sensitive to *SD* drive. The test *k* value (blue box) confirms this.  The orange box represents the reciprocal/control cross.

Fig S6

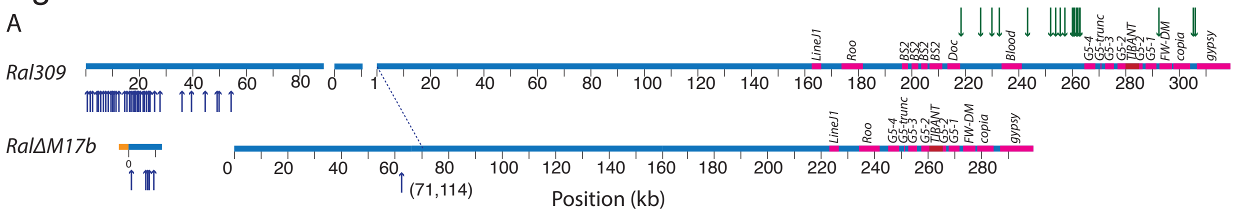


B

| | relative size semi-PCR (qPCR) | k value | # *Rsp* repeats sequence est. |
|---|---|---|---|
| *Ral309* | 1 | 0.97 | >2861 |
| *RalΔM5* (intermediate) | 0.53/0.37 (1.2/1.0/0.45) | 0.9 | nd |
| *RalΔM17b* | 0.93*/0.56/0.51 | 0.85 | ~2274 |

C

| | # of guide target sites | | |
|---|---|---|---|
| | start | post *Rsp-g1* CRISPR | post *Rsp-g2* CRISPR |
| *Rsp-g1* | | | |
| *Iso1ΔC16* | 31 | 3 | - |
| *Iso1ΔC8* | 31 | 0 | - |
| *Iso1ΔC16Δ20* | 3 | - | 3 |
| *Rsp-g2* | | | |
| *Iso1ΔC16* | 99 | 49‡ | - |
| *Iso1ΔC8* | 99 | 49‡ | - |
| *Iso1ΔC16Δ20* | 49 | - | 1 |
| *Rsp-g1* | | | |
| *RalΔM17b* | 19 | 0* | 0 |
| *Rsp-g2* | | | |
| *RalΔM17b* | 39 | 39* | 6 |

**Figure S6- Summary of target sites and sequential CRISPR in *Ral309*.**
A. Sequential CRISPR was performed on the *Rsp* array of the wildtype strain *Ral309.* In the first step, *Ral309* was subjected to editing with Cas9 and *Rsp-g1* (Figure S4), resulting in *RalM5* subline. *RalM5* was then subjected to editing with Cas9 and *Rsp-g2* (Figure S4), resulting in the subline *RalΔM17b*. The diagrams show the locations of *Rsp-g1* and *Rsp-g2* target sites and differences in non-*Rsp* sequences before and after CRISPR. B. Summarizes data from the PCR estimate assays, guide target site numbers, *k* values, and *Rsp* repeat numbers determined from sequencing. C. Compares the efficiency of target site cleavage between the *Iso1* and *Ral309* strains with the same guides. For each strain, the number of guide sites present at the start and post-CRISPR are shown. Dashes indicate the CRISPR cross was not done with that strain; double daggers indicate the missing 50 guide 2 target sites were involved in the large deletion generated with *Rsp-g1*; asterisks indicate assumed values.

Fig S7

A. Estimated *Rsp* counts

| Strain | *Rsp* Type | Count | Total Counts |
|---|---|---|---|
| *Iso1* | L-Rsp | 611 | 1756 |
| | R-Rsp | 1022 | |
| | Rsp-Variant | 50 | |
| | Rsp-Truncated | 73 | |
| *Iso1ΔC8* | L-Rsp | 300 | 851 |
| | R-Rsp | 500 | |
| | Rsp-Variant | 17 | |
| | Rsp-Truncated | 34 | |
| *Iso1ΔC16* | L-Rsp | 294 | 812 |
| | R-Rsp | 467 | |
| | Rsp-Variant | 17 | |
| | Rsp-Truncated | 34 | |
| *Iso1ΔC16Δ20* | L-Rsp | 57 | 211 |
| | R-Rsp | 114 | |
| | Rsp-Variant | 13 | |
| | Rsp-Truncated | 27 | |
| *Iso1ExpC11* | L-Rsp | 1233 | 3170 |
| | R-Rsp | 1642 | |
| | Rsp-Variant | 114 | |
| | Rsp-Truncated | 181 | |
| *Ral309* | L-Rsp | 1094 | 2861 |
| | R-Rsp | 1539 | |
| | Rsp-Variant | 102 | |
| | Rsp-Truncated | 135 | |
| *RalΔM17b* | L-Rsp | 895 | 2274 |
| | R-Rsp | 1169 | |
| | Rsp-Variant | 70 | |
| | Rsp-Truncated | 140 | |

B. *Rsp* locus in *Iso1* sublines

C. *Rsp* locus in *Ral309* sublines

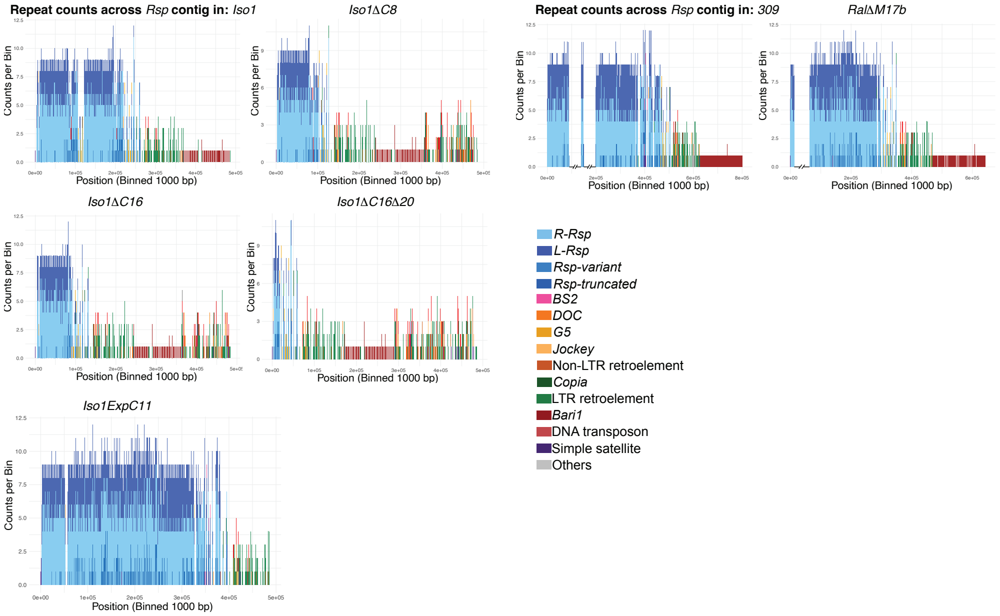


**Figure S7. *Rsp* copy number and organization.**
A. Table showing the total *Rsp* copy numbers and their different types based on their sequence length. For each subline, the barplots show the detailed structure of *Rsp* locus (see key) for B. *Iso1* sublines *Iso1ΔC16, Iso1ΔC8, Iso1ΔC16Δ20*, and *IsoExpC11;* and C. *Ral309* and its subline *RalΔM17b. Ral309* and *RalΔM17b* represent superscaffolds based on synteny patterns (see Fig S9).

Fig S8

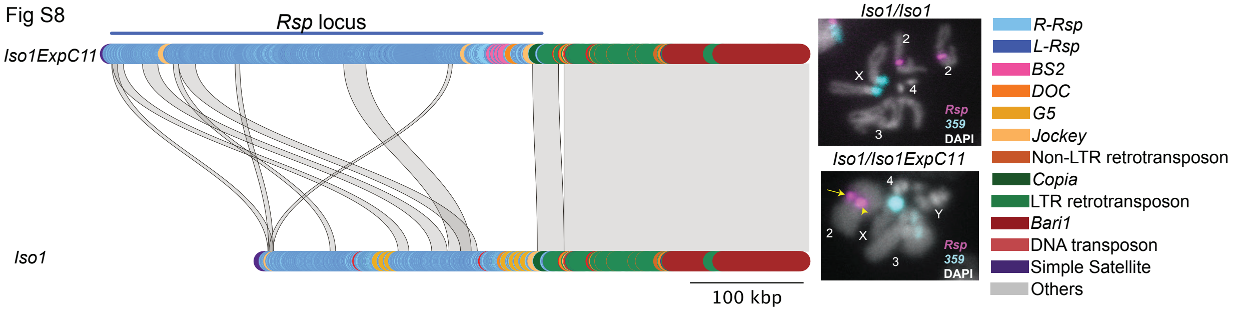


**Figure S8. Synteny plots for *Iso1* and *Iso1ExpC11*.**
*Iso1ExpC11* is identical to *Rsp* in the distal flanking region but the organization differs in *Rsp* organization, suggesting that it may originate from recombination between *Iso1* and the 2$^{nd}$ chromosome from the guide strain. Fluorescence in situ hybridization with *Rsp* probes and a control satellite not targeted by CRISPR (*359-bp*) confirm that the *Rsp* locus is expanded in *Iso1ExpC11.*

Fig S9

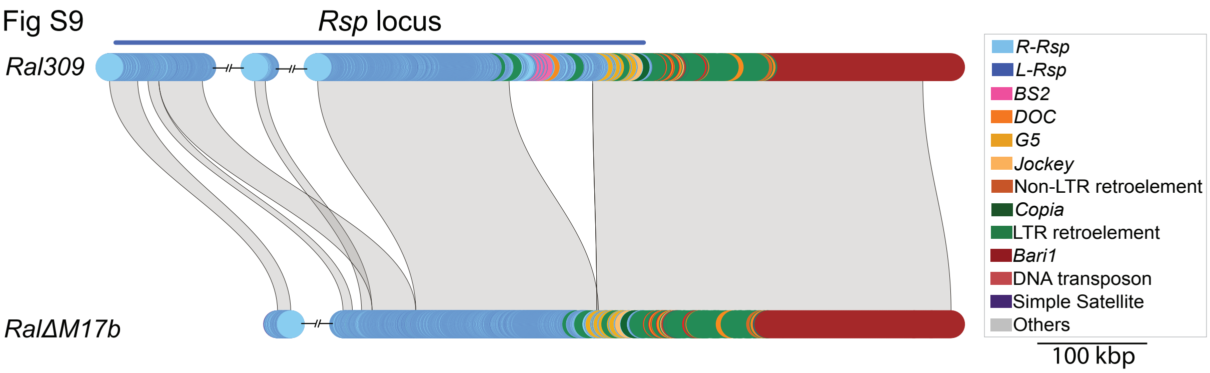


**Figure S9. Synteny plots for *Ral309* and *RalΔM17b*.**
The structure of the *Rsp* locus in *Ral309* differs from *Iso1* in *Rsp* copy number and organization, and in different interspersed TEs. The *RalΔM17b* has a deletions of *Rsp* and interspersed TEs on the distal side of the array. Note that because we did not get contiguous assemblies for Ral309 (total *Rsp* copy number 2861) and *RalΔM17b* (total *Rsp* copy number 2274), we created super scaffolds for each based on synteny.

Fig S10

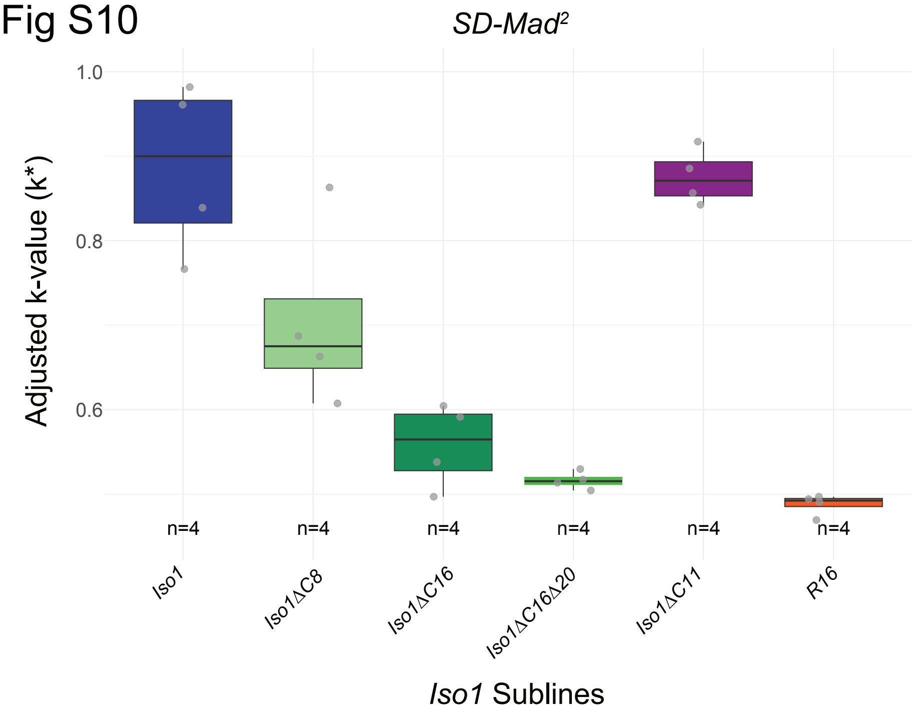


**Figure S10. Drive strength from intermediate driver of *SD-Mad$^2$* driver**
*SD-Mad* drive sensitivity decreases with reduced *Rsp* copy number. The difference in drive strength between *Iso1ΔC8* and *Iso1ΔC16* can be attributed to differences in *Rsp* copy number between the two sublines, with *Iso1ΔC16* having fewer *Rsp* repeats. Our control, *R16*—an X-ray mediated deletion of the pericentromeric region that completely lacks *Rsp*—shows no evidence of drive (FET P-value = 0.855; compared to *k* =0.5). The *Iso1ΔC16Δ20* subline which harbors a large *Rsp* deletion generated by sequential CRISPR editing of *Iso1ΔC16,* is also insensitive to *SD-Mad$^2$* (FET P-value = 0.119 compared to k = 0.5). *Iso1* and *Iso1ExpC11* show variation in k-values that are not strictly consistent with *Rsp* copy number, likely reflecting variability in *SD-Mad2* stability and genetic background effects (we may have introduced a suppressor when

replacing the genetic background). Nevertheless, the high k-values observed for *Iso1* (k = 0.887) and *Iso1ExpC11* (k = 0.875) indicate strong drive and are consistent with the presence of high *Rsp* copy numbers.

#### Supplemental Tables

Table S1

| guide name: sequence | # guide + PAM sites (# seed + PAM sites) | | off targets |
|---|---|---|---|
| *Rsp-g1*: GTACCCAAAAATAATTTGAATGG | 31(31) | | 2R: thisbe (ths)- fibroblast growth factor etc.<br>3R: Myosin 81F (Myo81F)- Myosin ATPase<br>3R: CG42342- extracellular matrix<br>X: kin of irre (kirre)- Myoblast aggregation etc.<br>2R: spaghetti-squash acxxx(sqa) myosin light chain kinase-like<br>3R: thisbe cherub- lncRNA:cherub<br>2R: pou domain motif 3 (pdm3)- odor receptor<br>3R: thisbe cherub- lncRNA:cherub<br>2L: CG10019- monocarboxylic acid transport<br>2L: Centaurin gamma 1A (CenG1A)- GTPase |
| *Rsp-g2*: GTTTGTCTGGTTCTTGAAAT | 99 (101) | 4 sites | 2L: 30C1; 3L: 76E2; 3R: 84A6<br>2L: Myosin 28B1 (Myo28B1)- high duty ratio motor protein<br>2L: Centaurin gamma 1A (CenG1A)- GTPase<br>3L: Argonaut 3 (Ago3)- piRNA guided cleavage activity |
| *Rsp-g3*: AATTTCCGATTTCAAGTACC | 121 (645) | 6 sites | 3L: Argonaut 3 (Ago3)- piRNA guided cleavage activity<br>3R: Myosin 81F (Myo81F)- Myosin ATPase<br>2R:: Dpr-interactlambda (DIP-lambda)- synaptic specificity<br>2R: Minor RSP locus<br>3R: NaKCl cotransporter (NKCC)- Na K Cl symporter activity<br>X: 4D7- repeat region |
| *Rsp-g4*: AGTTGAACAGAATCTCTAGA | 506 (539) | *4 sites | 3L: Argonaut 3 (Ago3)- piRNA guided cleavage activity<br>3R: CG14891- nucleic acid binding etc.<br>3L: Argonaut 3 (Ago3)- piRNA guided cleavage activity<br>3L: Argonaut 3 (Ago3)- piRNA guided cleavage activity |

**Table S1: Predicted number of guide targets and off targets**
The guide name and sequence for each of the 4 guide RNAs used is shown in the first column. The number of perfect target guide sites is presented in the second column, followed by the number of perfect seed sites in parentheses. Potential off-target sites outside of the *Rsp* array as predicted by flyCRISPR are listed in the last column. None of the potential targets outside of the *Rsp* array were perfect, with the exception of the 4 sites (marked with asterisk) in the Ago3 gene which is a perfect target for *Rsp-g4*.

Table S2

| allele_original name | estimated # of *Rsp* repeats | | | | k value |
|---|---|---|---|---|---|
| | qPCR | slot blot | PCR <2019 | PCR 2019 | (*Iso1*=0.91) |
| *IsoCΔ1_F22-1FG* | 278/202 | | 874 | 1269 | |
| *IsoCΔ2_F22-2FC* | 26/135 | 1486/1012 | | 1186 | 0.843 |
| *IsoCΔ3_F22-3FG* | 234/641 | | | 1137 | |
| *IsoCΔ4_F22-4FC* | 30/241 | | 1046 | 795 | |
| *IsoCΔ5_F22-5FG(CyO)* | 2174/229 | | | 2x239 | |
| *IsoCΔ6_F22-6FC* | 556/248 | | | 1173 | |
| *IsoCΔ7_M14-3FC(CYO)* | 1780/1634 | 2183/1624 | 1063 | 2x421 | |
| *IsoCΔ8_M14-8MG* | 359/397 | 586/613 | 709 | *122* | 0.686 |
| *IsoCΔ9_M2A-M3C(CYO)* | | | (650) | 2x46 | |
| *IsoCΔ10_M2A-M4C(CyO)* | | | | 2x1398 | |
| *IsoCΔ11_M2A-M4C* | | | 1800 | 1874 | 0.947 |
| *IsoCΔ12_M7H-M10G(Cy0)* | | | 2x1100 | 2x638 | |
| *IsoCΔ13_F3D-M3Cr(CyO)* | | | | 2x443 | |
| *IsoCΔ14_F3D-M3Cr* | | | 564 | 1167 | |
| *IsoCΔ15_F3D-M5Cv(CyO)* | | | | 2x111 | |
| *IsoCΔ16_F3D-M5Cv* | | | 533 | 484 | 0.646 |
| *IsoCΔ17_F8A-M4Cwy(CyO)* | | | | <100* | |
| *IsoCΔ18_F8A-M4Cwy* | | | 587 | 382 | 0.82 |
| *IsoCΔ19_F8A-M5Cwy(CyO)* | | | | 2x279 | |
| *IsoCΔ20_F8A-M5Cwy* | | | 695 | 372* | |
| *IsoCΔ21_F15E-M3Gv(CyO)* | | | (250) | 2x73* | |

**Table S2. Assessment of *Rsp* alleles.**
Twenty one strains generated in the original CRISPR crosses (Figure 1) were selected for further analysis. A second set of semi-quantification PCRs and/or slot blots were performed. Those strains which had consistent *Rsp* repeat estimates were assayed for their sensitivity to *SD* distortion (*k* value). Strains *IsoΔC8*, *IsoΔC11*, and I*soΔC16* (highlighted in green) were selected for additional analysis.

Table S3

A

| Strain | Choromosome arm | Corresponding contig in *Iso1* | Est. recombination interval | Number of SNPs outside breakpoints |
|---|---|---|---|---|
| *Iso1ΔC8* | 2L | ptg000025l | NA | 7 |
| | 2L | ptg000075l | NA | 6 |
| | 2R | ptg000018l | NA | 6 |
| | 2R | ptg000023l | NA | 6 |
| | 2R | ptg000047l | NA | 1 |
| *Iso1ΔC16* | 2L | ptg000025l | 1056922, 12726395 | 15 |
| | 2L | ptg000075l | NA | 4 |
| | 2R | ptg000018l | NA | 2 |
| | 2R | ptg000023l | NA | 5 |
| | 2R | ptg000047l | NA | 4 |
| *Iso1ΔC16Δ20* | 2L | ptg000025l | 1056922, 12490127 | 12 |
| | 2L | ptg000075l | NA | 6 |
| | 2R | ptg000018l | NA | 6 |
| | 2R | ptg000023l | NA | 6 |
| | 2R | ptg000047l | NA | 2 |
| *Iso1ΔC11* | 2L | ptg000025l | 1050210, 21472445 (END) | 12<br>6 |
| | 2L | ptg000075l | 301198, 2022672 (END) | 6<br>6 |
| | 2R | ptg000018l | NA | 2 |
| | 2R | ptg000023l | NA | |
| | 2R | ptg000047l | NA | |

B

| Strain | Choromosome arm | Corresponding contig in *Ral309* | Est. recombination interval | Number of SNPs outside breakpoints |
|---|---|---|---|---|
| *RalΔM17b* | 2L | ptg000029l | NA | 0 |
| | 2L | ptg000015l | NA | 2 |
| | 2L | ptg000063l | NA | 0 |
| | 2R | ptg000004l | NA | 3 |

**Table S3. SNPs and putative recombination events**

We summarize putative recombination event and mutations that accumulate on the 2nd chromosomes (not involving the recombination events) for *Iso1* sublines (A) and *RalΔM17b* (B). *Iso1ΔC8* did not have any recombination event on chromosome 2, consistent with its generation using injection-based scheme. In contrast, *Iso1ΔC16* and *Iso1ΔC16Δ20* show a putative recombination breakpoint at similar positions along the 2L chromosome arm (presumably the same event since *Iso1ΔC16Δ20* is derived from *Iso1ΔC16*), while *IsoExpC11* has presumed recombination intervals spanning most of chromosome 2L. We do not detect any evidence of recombination on chromosome 2R. We also report the number of variant sites outside of the inferred recombination regions on chromosome 2L and 2R. Consistent with the timeline of line generation, these variants are unlikely to have arisen from the CRISPR event itself and instead likely represent the accumulation of mutations since the origin of each line. *RalΔM17b* does not exhibit any recombination events, and consistent with the recent generation from the parental *Ral309* strain, shows a reduced number of variants on chromosome 2.
